# FAMeDB: A curated Database for the analysis of Fungal Aromatic Compound Metabolism

**DOI:** 10.64898/2026.01.19.700319

**Authors:** Tiago M. Martins, Rita C. Carmo, Cristina Silva Pereira

## Abstract

Aromatic compounds represent the second most abundant class of organic molecules after carbohydrates, and their microbial metabolism is of broad relevance across multiple research disciplines. Metabolic pathways involving aromatic compounds span from highly conserved anabolic routes to more variable catabolic processes. Numerous peripheral catabolic pathways converge on a small number of central intermediates that undergo aromatic ring-opening in the central pathways. Over the past decades, alongside numerous peripheral pathway genes, most of the catabolic genes constituting the central metabolic pathways have finally been characterized in fungi. Here we present FAMeDB, a manually curated database of proteins involved in fungal aromatic compound metabolism, together with associated bioinformatic tools. The database currently includes 349 proteins, primarily enzymes, but also includes transcription factors and transporters. Entries span 65 species and 45 genera of fungi. Most entries are from Ascomycota (81%), with a significant number from Aspergilli (46%). Application of FAMeDB and its tools enables the quick and accurate representation of aromatic metabolism across different fungal proteomes. This resource is designed to provide a useful and accessible platform for researchers worldwide, even those without specialized expertise in fungal aromatic catabolism, facilitating omics analysis and genomic comparisons.

**GRAPHICAL ABSTRACT:** 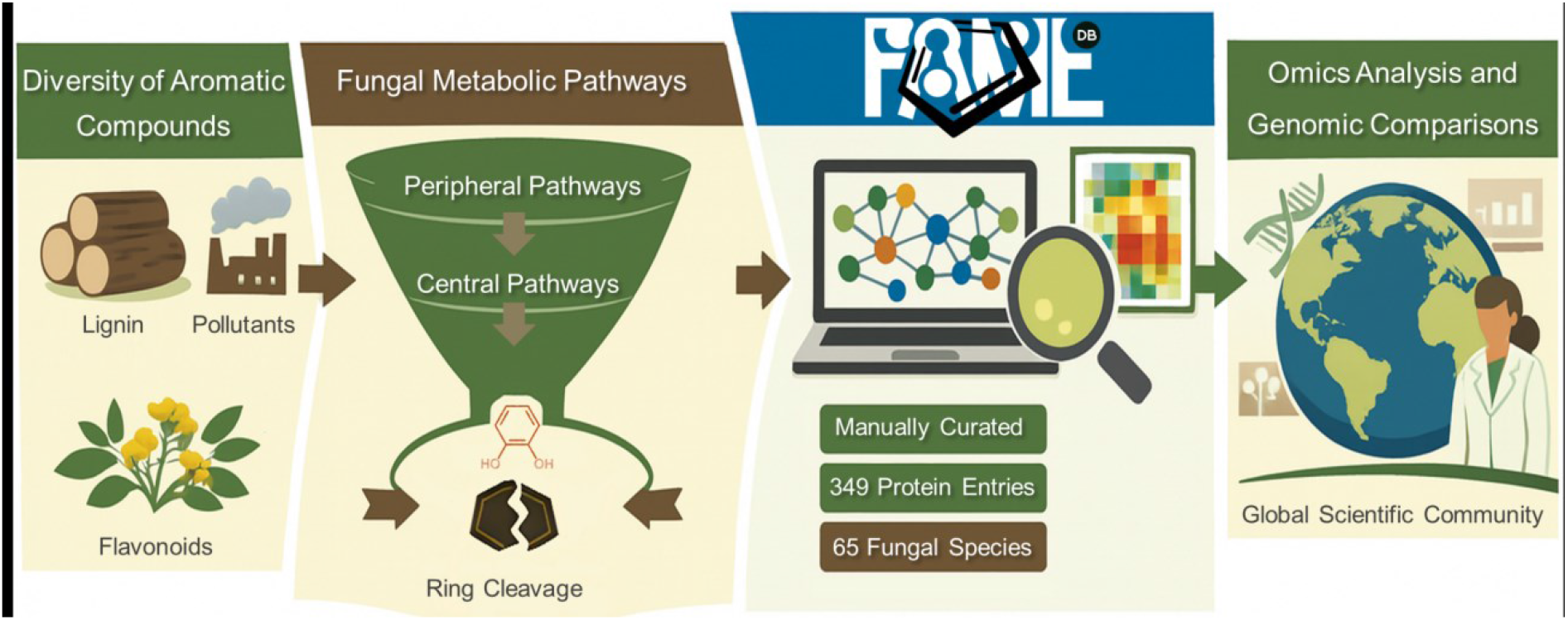

## INTRODUCTION

Aromatic compounds constitute the second most abundant class of organic molecules in nature, surpassed only by carbohydrates. Their structural diversity and chemical stability underpin a wide array of biological functions and ecological interactions, making their metabolism a focal point across disciplines ranging from environmental microbiology to industrial biotechnology. Microbial degradation and transformation of aromatics are particularly critical for nutrient cycling, bioremediation, and the biosynthesis of high-value metabolites. Despite extensive research in bacterial systems, fungal aromatic metabolism remains comparatively underexplored, even though fungi play pivotal roles in terrestrial ecosystems and industrial processes.

Fungi exhibit distinct metabolic strategies that set them apart from other microbial groups, particularly in their ability to process structurally complex aromatic compounds. Fungi contribute significantly to lignin degradation and the turnover of plant-derived aromatic polymers, processes essential for carbon cycling and soil health (1). Recent advances in fungal research in the omics era have revealed a wealth of previously uncharacterized genes and pathways involved in aromatic compound metabolism (2–6). To date, eight central ring-opening intermediates in the catabolism of aromatic compounds have been identified in fungi: 1,2,3,5-tetrahydroxybenzene, 3-hydroxyanthranilate, catechol, gentisate, homogentisate, hydroxyquinol, methoxyhydroquinone, and protocatechuate. This surge in data, coupled with the increasing availability of fungal genomes has opened new avenues for comparative and functional analyses, yet the lack of centralized, curated resources has hindered systematic exploration of fungal aromatic metabolism.

Existing resources relevant to the analysis of fungal aromatic compound catabolism include CAZy (7), HADEG (8), KEGG (9), and BioCyc (10). However, their utility for investigating fungal metabolic pathways involving aromatic compounds remains limited. For instance, CAZy primarily catalogs carbohydrate-active enzymes, with only a subset of auxiliary enzymes (*e*.*g*., those acting on lignin) implicated in aromatic compound degradation. Other databases tend to be either incomplete or predominantly focused on non-fungal taxa (*e*.*g*., HADEG). Moreover, key biochemical differences between fungal and bacterial central catabolic pathways (*e*.*g*., involving 3-carboxymuconolactone and its associated enzymes) are often overlooked (*e*.*g*., KEGG and BioCyc), even if being well-documented for decades.

To address current gaps and complement existing resources, we present FAMeDB - a curated database of protein entries dedicated to the fungal metabolism of aromatic compounds. Alongside this, we include bioinformatic tools designed to facilitate rapid characterization of fungal proteomes regarding their metabolic potential in response to aromatic substrates.

## MATERIAL AND METHODS

### Sequence collection

Proteins were mostly identified through literature curation. Additional orthologs with protein-level evidence in UNIPROT were obtained via BLAST searches against the curated SWISS-PROT database, using initially identified entries as queries. Protein sequences and associated descriptive data were retrieved from public repositories, including UNIPROT (11), NCBI (12), and Mycocosm (13). Complementary annotation of protein entries was performed using eggNOG-mapper (14) and InterProScan (15). Carbohydrate active enzymes (CAZymes) are not the object of this database and were removed after dbCAN3 search (16), except for the dual tannase / beta-glucosidase BglA and vanillyl-alcohol oxidase VaoA.

### Orthology and database analysis

Proteinortho v. 6.3.6 (17) was used to detect orthologous genes between database entries and different query species proteomes. Default parameters were used except algebraic connectivity was set at 0.3 and identity was set at 40 or at different values as otherwise indicated. Analysis of fungal proteomes with HADEG database was also performed as previously described (8). For comparison purposes with FAMeDB, the HADEG database was also used directly in orthology analysis. To analyze 150 fungal proteomes, we first selected reference annotated genomes from NCBI (<10,000 assembled scaffolds) containing between 5,000 and 30,000 coding genes. From this set, we randomly selected 50 genomes from each of the Ascomycota, Basidiomycota, and non-Dikarya lineages (for accession numbers see Table S1).

Orthology data were analyzed in R (v. 4.5.1) using the packages stringr (v. 1.5.1), dplyr (v. 1.1.4), readr (v. 2.1.5), scales (v. 1.4.0), svglite (v. 2.2.1), ggplot2 (v. 3.5.2), and reshape2 (v. 1.4.4) to identify database-matching hits by pathway and to generate the corresponding tables and plots.

## RESULTS

The FAMeDB curated database currently includes 349 proteins (231 orthology groups), mostly enzymes but also some transcription factors and transporters. The respective pathways range from peripheral catabolism to central metabolism, including some that do not directly deal with aromatic compounds but may be upstream or downstream of them (*e*.*g*., quinic acid metabolism) or that belong to secondary metabolism. The pathways include anabolic and catabolic pathways, as it is often impossible to discern molecular components from each other, as they encompass redundancy and plasticity, even more so when considering secondary metabolism.

The database contains entries from 65 species and 45 genera of fungi. Around half are from Aspergilli (46%), and the vast majority from Ascomycota (81%). This distribution primarily reflects the fungal species most frequently employed in research, particularly those studied in the context of aromatic compound metabolism.

Among existing resources, the HADEG database offers coverage of aromatic compound metabolism (8). However, its utility for fungal systems is limited: only three fungal entries are included (although not in aromatics metabolism), and overall orthology and homology between bacterial and fungal enzymes is low. An analysis of 150 random fungal proteomes using the HADEG database yielded only a few specific matches primarily for homogentisate 1,2-dioxygenase HmgA and catechol 1,2-dioxygenase CatA (Figure S1 and Table S2). The analysis also revealed some non-specific or unknown hits, including maleylpyruvate isomerase NagL (shows homology to fungal maleylacetoacetate isomerase), 3-oxoadipyl-CoA thiolase PcaF (lacking orthology to characterized Ascomycota counterparts), and 5-carboxymethyl-2-hydroxymuconate semialdehyde dehydrogenase HpcC (homoprotocatechuate degradation, which is not characterized in fungi). The divergence to bacteria is especially pronounced in certain catabolic pathways, such as the fungal-specific 3-carboxymuconolactone route within the protocatechuate branch (2), or the (yet) unique fungal 1,2,3,5-tetrahydroxybenzene central pathway (6). Attempts to circumvent this limitation by lowering identity thresholds to detect distant homologs often compromise specificity, increasing the likelihood of false positives - proteins that are structurally related but functionally divergent (*e*.*g*., superfamily members). A problem that is significantly reduced when using FAMeDB for the analysis of fungal proteomes. Nevertheless, this methodological limitation should be considered when analyzing species outside Ascomycota, given the database’s current bias toward this phylum.

Curation of FAMeDB prioritized entries with robust experimental characterization. Nonetheless, proteins with limited evidence - such as those within conserved gene clusters - were included to encourage further investigation and functional validation. While the primary goal was to facilitate identification of the more divergent catabolic pathways, comprehensive annotation of related metabolic pathways was necessary to ensure contextual accuracy.

Certain enzyme families, such as CAZymes involved in peripheral degradation (*e*.*g*., lignin peroxidases), are better accessed through specialized resources like CAZy DB and dbCAN3 (16, 18). Contrarily, conserved biosynthetic pathways (*e*.*g*., coenzyme Q10 synthesis) were retained in the database, as their constituent genes may have been repurposed for novel functions. These distant paralogs are particularly prone to misannotation, especially when comparing them across distantly related taxa.

Another conserved pathway - the nicotinate metabolism - was also included due to its potential intersection with aromatic catabolism via the upper 3-hydroxyanthranilate pathway, which may compensate for the absence of the lower pathway in certain species such as those of genus *Candida* (5, 19). Although further research is needed, its relevance warrants inclusion. On the other hand, Basidiomycota generally retains NAD biosynthesis and salvage genes but lack many of the nicotinate degradation genes (20).

Pathway annotation also required careful consideration of phylum-specific differences. For instance, Basidiomycota variants (different gene architecture or orthology) of the catechol and protocatechuate branch in the 3-oxoadipate pathway are supported primarily by conserved gene cluster evidence, including protocatechuate dioxygenase (4). These differences, though preliminary, should not be overlooked. Despite the genetic differences, they were annotated in the corresponding pathway as there is no evidence of biochemical distinctions between fungal phyla nor is there evidence that both gene compositions could be present in a single organism.

The gentisate-like pathway presents additional challenges. It lacks a maleylpyruvate isomerase, and its functional role remains unclear despite genomic co-occurrence with the canonical gentisate pathway (4, 21). Transcriptomic data indicates upregulation in gentisate- or lignin-containing media (22), but definitive functional validation is pending. Notably, Basidiomycota gentisate-like clusters also omit this isomerase, suggesting it may be non-essential or involved in detoxification of metabolites of gentisate derivatives. Distinct catalytic properties of gentisate 1,2-dioxygenases from these clusters (23) support the hypothesis of isoenzyme specialization. Nevertheless, the distinction was only made in the annotation of the peripheral pathways, as the central pathway proteins are conserved and cannot be distinguished by the current methodology.

Functional or enzymatic redundancy is another important consideration. For example, maleylacetate reductase plays roles in both hydroxyquinol and methoxyhydroquinone pathways (24, 25). Gallate 1-monooxygenase (decarboxylating), can play a role upstream of the 1,2,3,5-tetrahydroxybenzene pathway or the hydroxyquinol branch, as it also can metabolize *e*.*g*. protocatechuate (6, 25, 26). The 3-carboxy-*cis,cis*-muconate cyclase paralog in Basidiomycota has a putative role as a *cis,cis*-muconate cycloisomerase even if joined together with 3-oxoadipate-enol lactonase in a multifunctional enzyme (4). Similarly, enzymes in CoA-dependent pathways, such as those in the 3-oxoadipate pathway or β-oxidative routes for hydroxycinnamic acids, are prone to produce false positives or misannotation using FAMeDB analysis due to high family-level redundancy. For instance, the high sequence homology between the orthologs 3-oxoadipyl-CoA thiolase KctA (3-oxoadipate pathway) and 3-ketoacyl CoA thiolase KatA (cinnamic acids CoA-dependent beta-oxidative pathway) often leads to misannotation, as the methodology described here lacks the necessary resolution to fully distinguish them, even within the same phyla.

These nuances in database construction should inform interpretation of results derived from FAMeDB. Nevertheless, application of the database and its associated tools to 150 randomly selected fungal genomes across multiple phyla (Figure 1 and Table S3) demonstrates its capacity to deliver rapid, informative insights into fungal aromatic metabolism.

**Figure 1.**
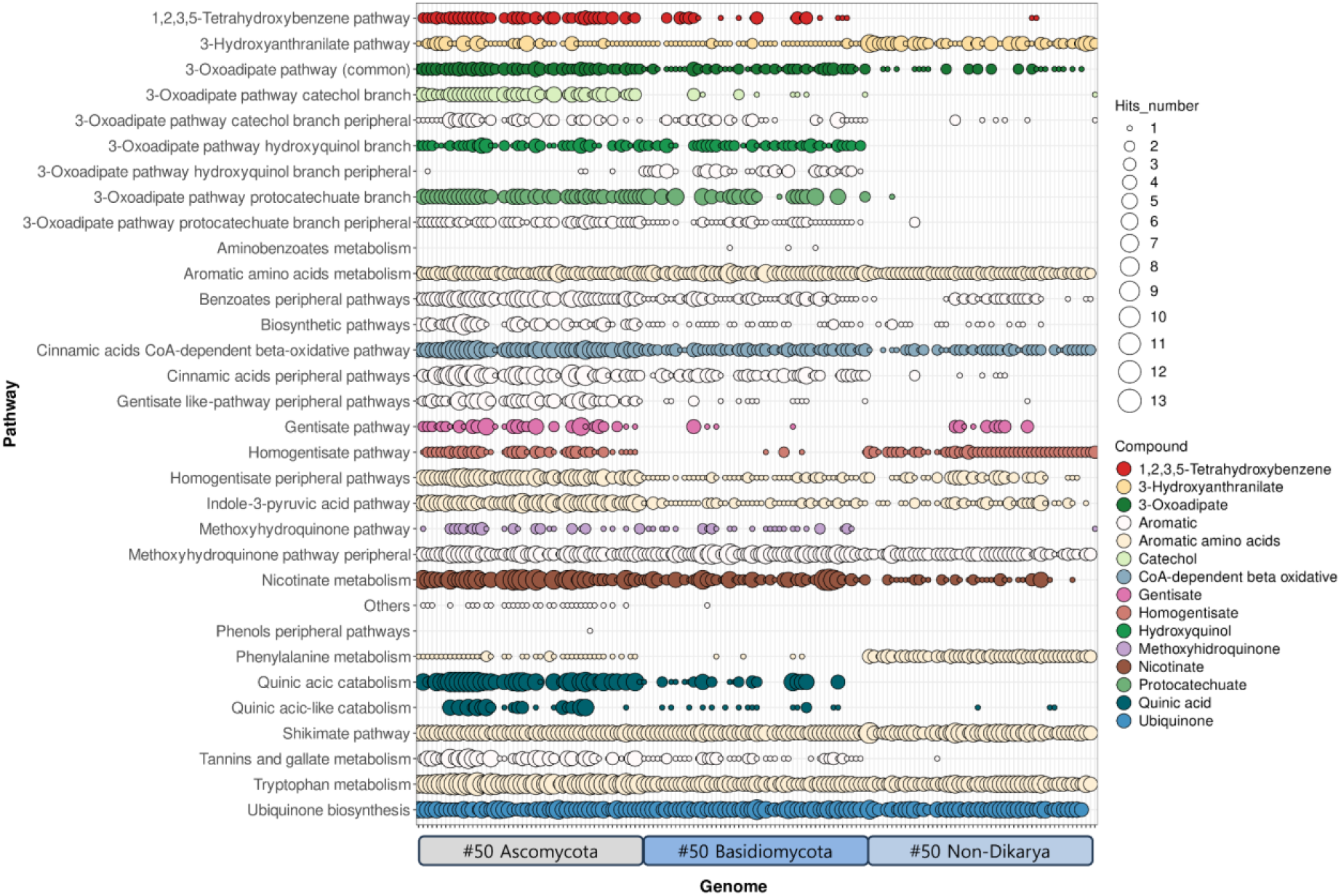
FAMeDB analysis of 150 random fungal proteomes. The number of database hits annotated per pathway is displayed using a bubble plot. Bubbles are colored according to compound. The rightmost column provides a direct comparison of matches obtained against HADEG database. Genomes are sorted alphabetically by species name in their corresponding taxonomic group (see Table S1). Results shown consider a minimum sequence identity of 40% in the orthology analysis step.

Overall, the analysis of the results obtained with 150 fungal proteomes (Figure 1 and S2 to S6, and Table S3 to S4) quickly indicates several key differences between the three taxonomic groups under analysis. Ascomycota genomes generally have a larger repertoire of central pathways, although caution should be exercised as more research has been conducted on this fungal taxon. Consequently, some proteins may not be properly identified for the other taxa in the current version of FAMeDB. It can also be observed that, in general, some central pathways are less frequently present in Basidiomycota species and that they have limited nicotinate metabolism. The last is better observed in a heatmap plot of the presence of the genes of this pathway (Figure 2), which shows that the HnxS nicotinate hydroxylase is less frequently present, as are the other genes, in Basidiomycota than in Ascomycota. Non-Dikarya fungi have a more limited set of catabolic central pathways - at least among those currently known (Figures 1, 3, and S2 to S6). Nonetheless, non-Dikarya fungi frequently possess the 3-hydroxyanthranilate and homogentisate pathways, which may be important for the metabolism of aromatic amino acids.

**Figure 2.**
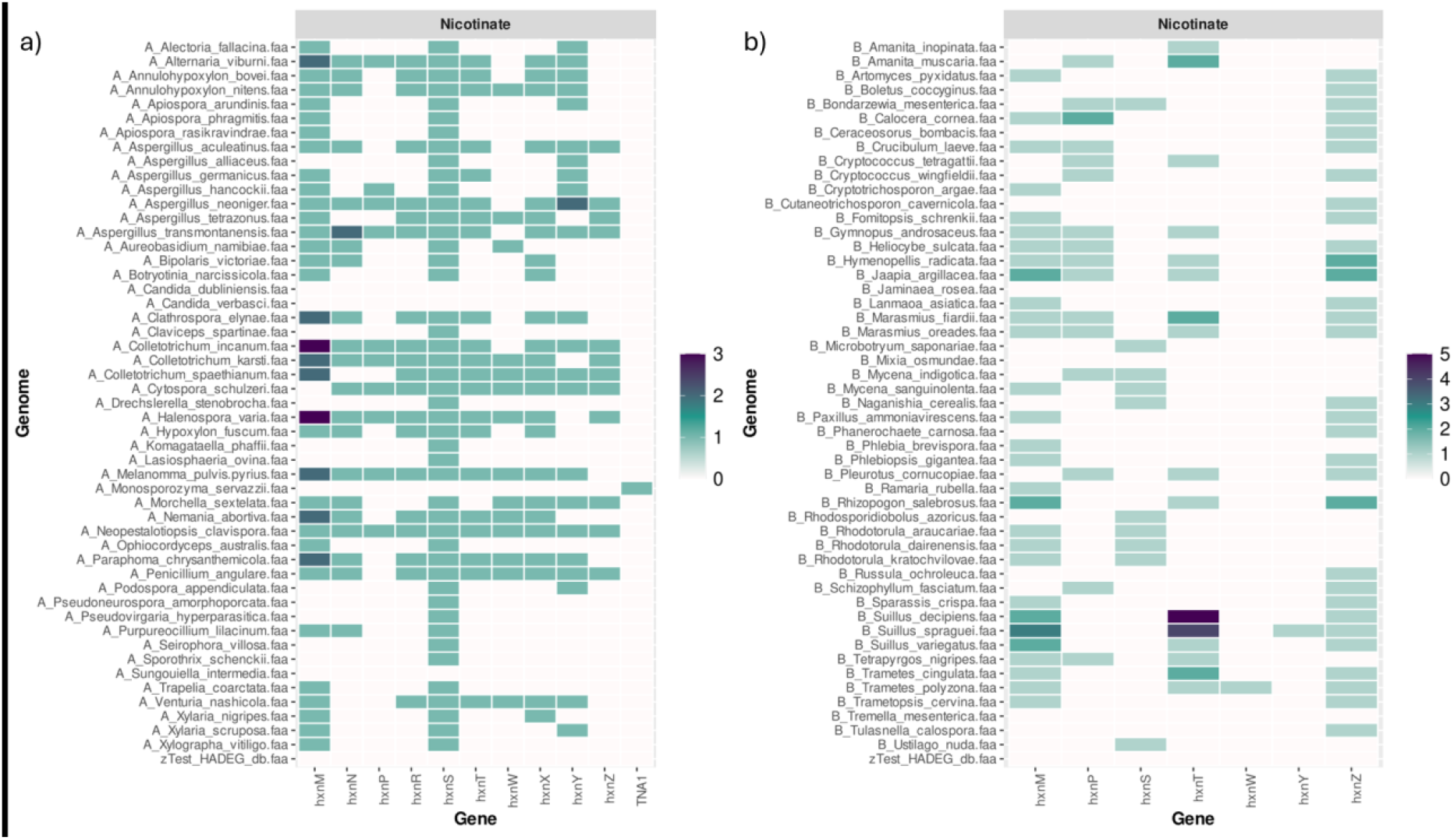
FAMeDB analysis of nicotinate metabolism in fifty random proteomes of Ascomycota (a) and fifty of Basidiomycota (b). Genes of nicotinate metabolism and in particular nicotinate hydroxylase HnxS are less frequently present in Basidiomycota in comparison to Ascomycota genomes. The number of database hits of the nicotinate metabolism annotated per proteome is displayed using a heatmap plot. The last row provides a direct comparison of matches obtained against HADEG database. Results shown consider a minimum sequence identity of 40% in the orthology analysis step.

**Figure 3.**
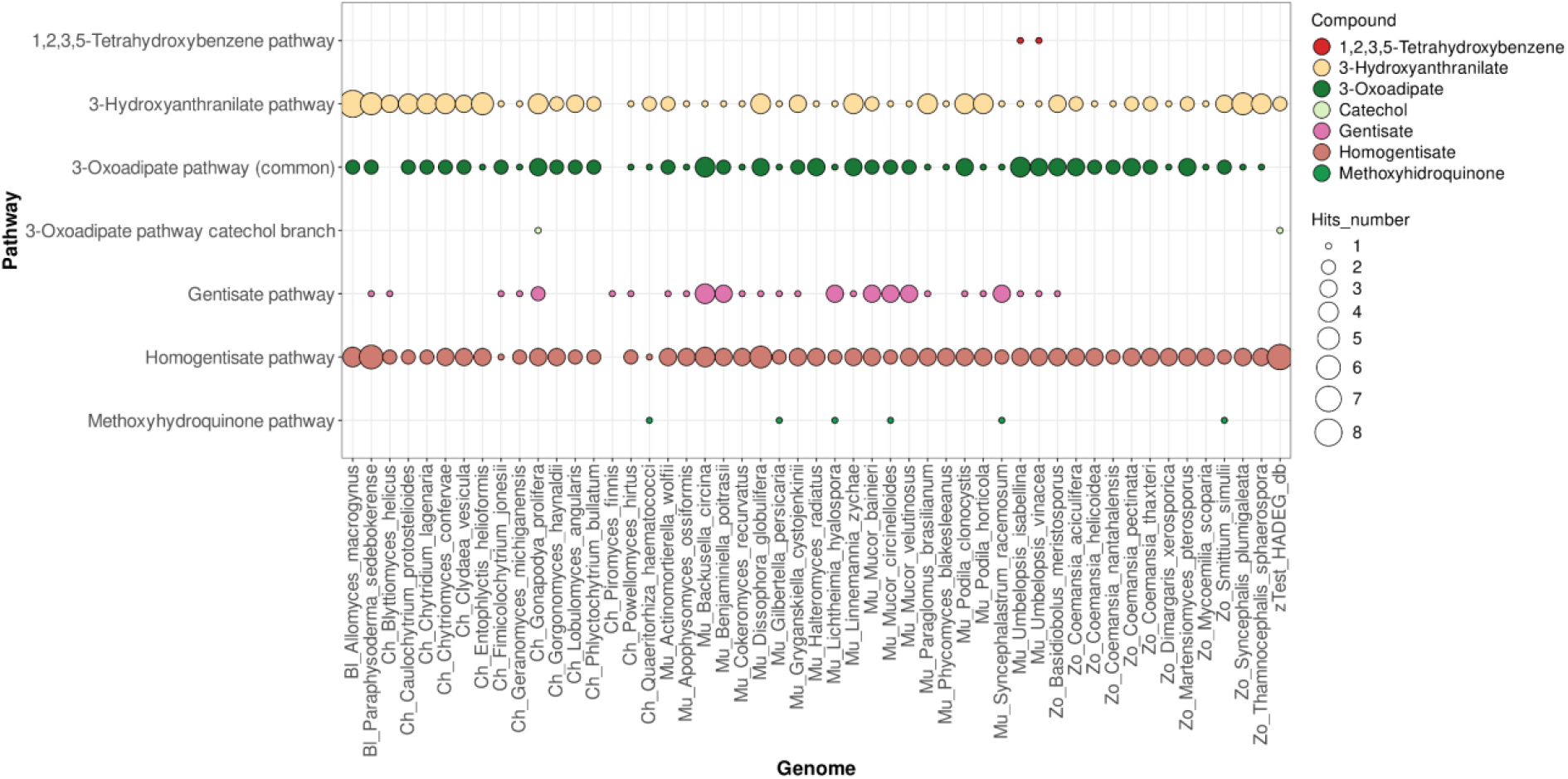
FAMeDB analysis of the central pathways for aromatic compounds catabolism of fifty random fungal proteomes of non-Dykaria. The number of database hits annotated per pathway is displayed using a bubble plot. Bubbles are colored according to compound. The rightmost column provides a direct comparison of matches obtained against HADEG database. Results shown consider a minimum sequence identity of 40% in the orthology analysis step.

Setting the identity at 40% in the orthology step allows a broader identification of hits and permits to withstand the wide differences between species in the fungi kingdom. However, the identification of a single hit or multiple hits for the same protein is common and reflects the expected redundancy. Therefore, the presence of a certain pathway should only be admitted if two or more different enzymes are identified.

## DISCUSSION

FAMeDB enables the rapid and informative analysis of fungal genomes, shedding light on their aromatic metabolism. Consequently, this toolset can be used to integrate knowledge and accelerate future research across various fields that study aromatic metabolism.

This database is inherently limited by the current understanding of fungal aromatic catabolism, including the input from its curators. To remain relevant and helpful, it should be constantly updated, preferably with the help of the research community.

The analysis methodology reported in this work is highly dependent on how well annotated the genomes under study are, or even if annotated at all at gene level. Absent or wrongly annotated genes (*e*.*g*., shorter) in a genome annotation can give the suggestion that a specific pathway may not be present. In a future development, this can be mitigated if the analysis can be performed on the genome sequence directly.

## ACKNOWLEDGEMENTS

We acknowledge André Cairrão for the design of FAMeDB logo.

## AUTHOR CONTRIBUTIONS

Tiago M. Martins (Conceptualization, Data curation, Formal analysis, Funding acquisition, Investigation, Methodology, Project administration, Resources, Software, Validation, Visualization, Writing – original draft, Writing – review & editing), Rita C. Carmo (Data curation) and Cristina Silva Pereira (Conceptualization, Funding acquisition, Project administration, Resources, Supervision, Writing – review & editing).

## DATA AVAILABILITY

All data associated with FAMeDB - including protein sequences and annotations, pathway descriptive table, and accompanying R scripts - is publicly available in the GitHub repository (https://github.com/SilvaPereiraLab/FAMeDB).

