## Supplementary Figures for "FAMeDB: A curated Database for the analysis of Fungal Aromatic Compound Metabolism"

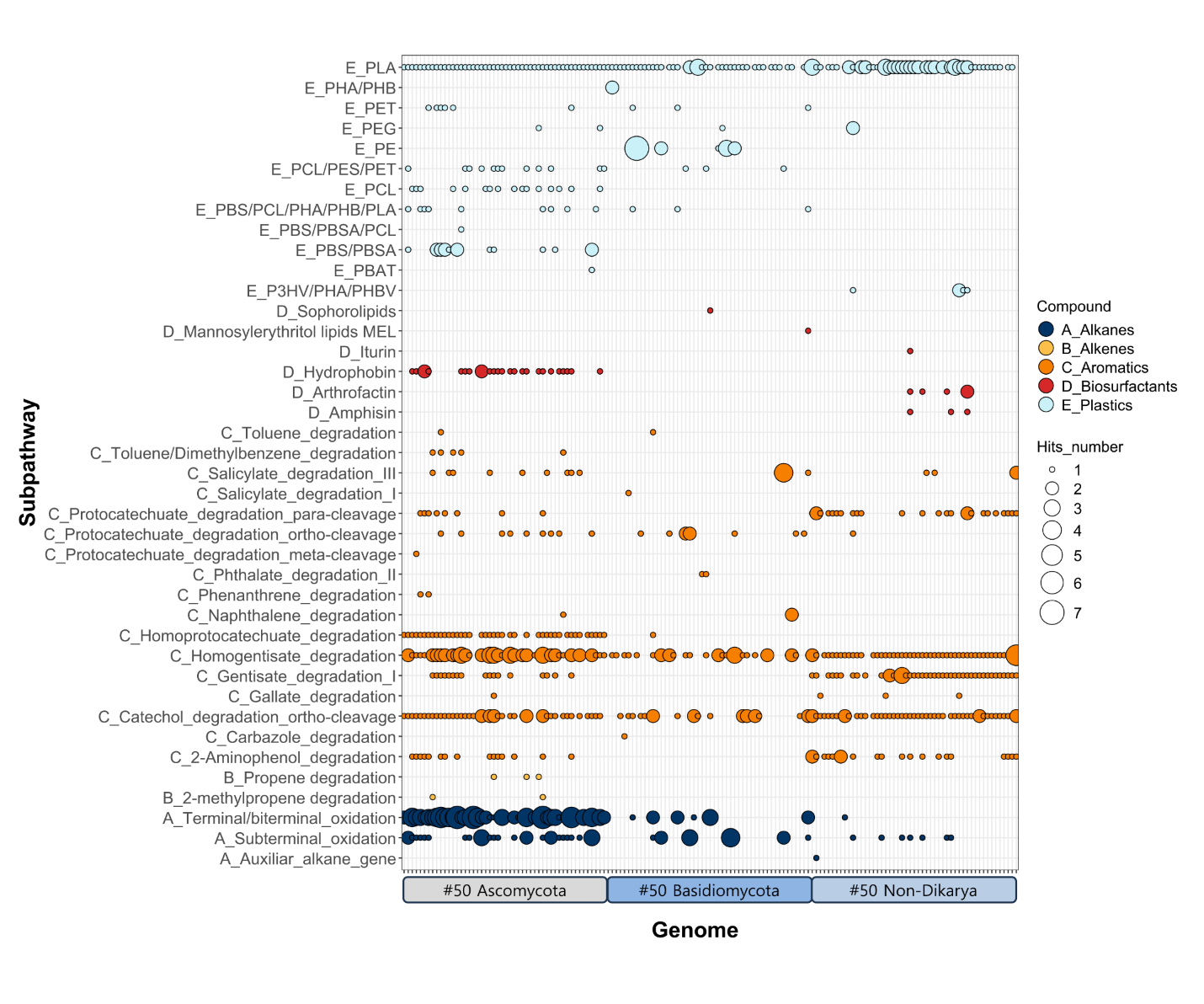

**Figure S1**. HADEG database analysis of 150 random fungal genomes. The number of database hits annotated per pathway is displayed using a bubble plot. The rightmost column provides a direct comparison of matches obtained against FAMeDB. Genomes are sorted alphabetically by species name in their corresponding taxonomic group (see Table S1). Results shown consider a minimum sequence identity of 40% in the orthology analysis step.

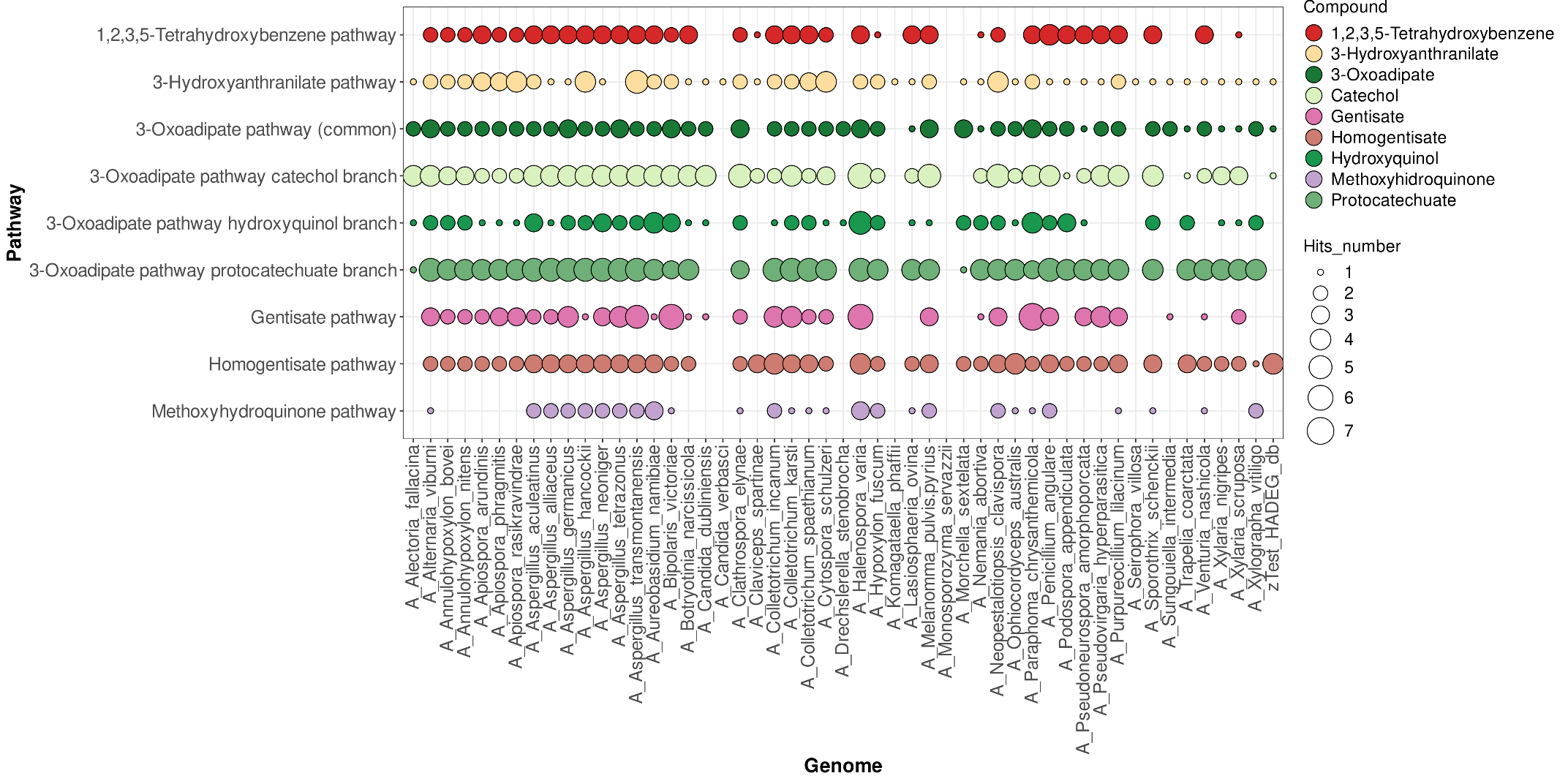

**Figure S2**. FAMeDB analysis of the central pathways for aromatic compounds catabolism of fifty random fungal proteomes of Ascomycota. The number of database hits annotated per pathway is displayed using a bubble plot. Bubbles are colored according to compound. The rightmost column provides a direct comparison of matches obtained against HADEG database. Results shown consider a minimum sequence identity of 40% in the orthology analysis step.

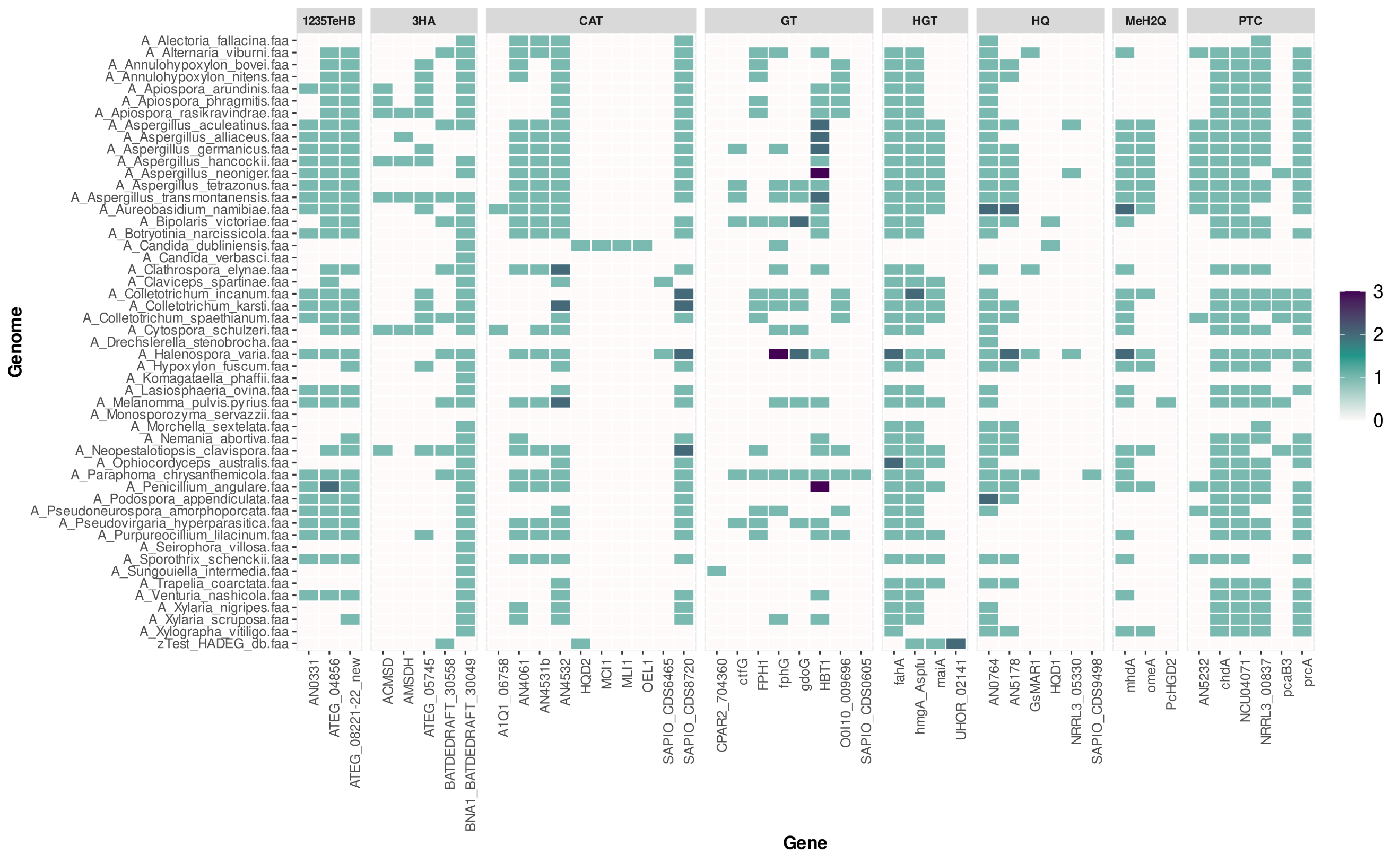

**Figure S3**. FAMeDB analysis of central pathways for aromatic compounds catabolism in fifty random proteomes of Ascomycota. The number of database hits of the central pathways annotated per proteome and gene (representing an ortholog group) is displayed using a heatmap plot. The last row provides a direct comparison of matches obtained against HADEG database. Results shown consider a minimum sequence identity of 40% in the orthology analysis step. Abbreviations: 1235TeHB - 1,2,3,5‑tetrahydroxybenzene, 3HAO - 3‑hydroxyanthranilate, CAT - catechol, GT - gentisate, HGT - homogentisate, HQ - hydroxyquinol, MeH2Q - methoxyhydroquinone, and PTC – protocatechuate.

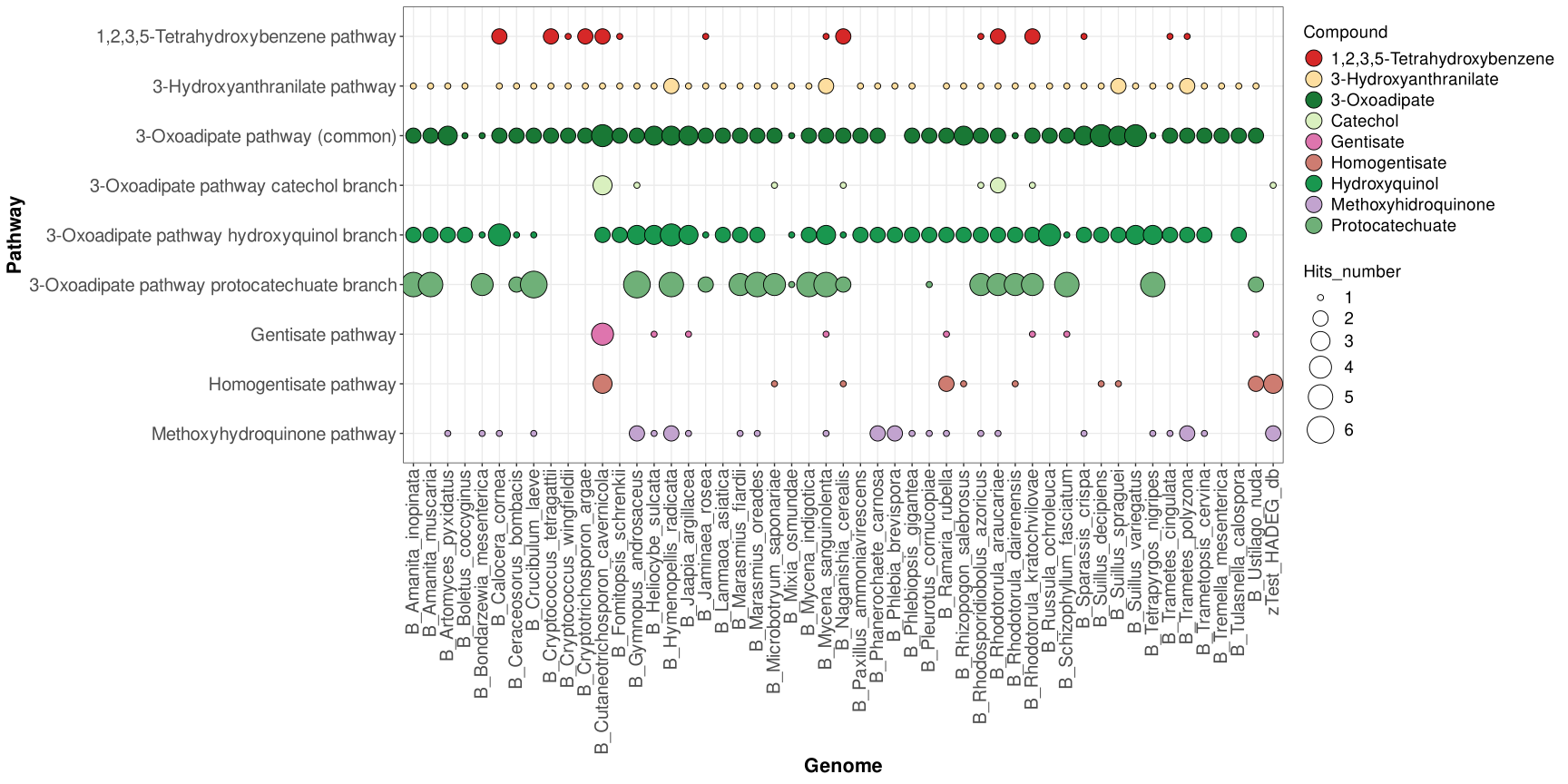

**Figure S4**. FAMeDB analysis of the central pathways for aromatic compounds catabolism of fifty random fungal proteomes of Basidiomycota. The number of database hits annotated per pathway is displayed using a bubble plot. Bubbles are colored according to compound. The rightmost column provides a direct comparison of matches obtained against HADEG database. Results shown consider a minimum sequence identity of 40% in the orthology analysis step.

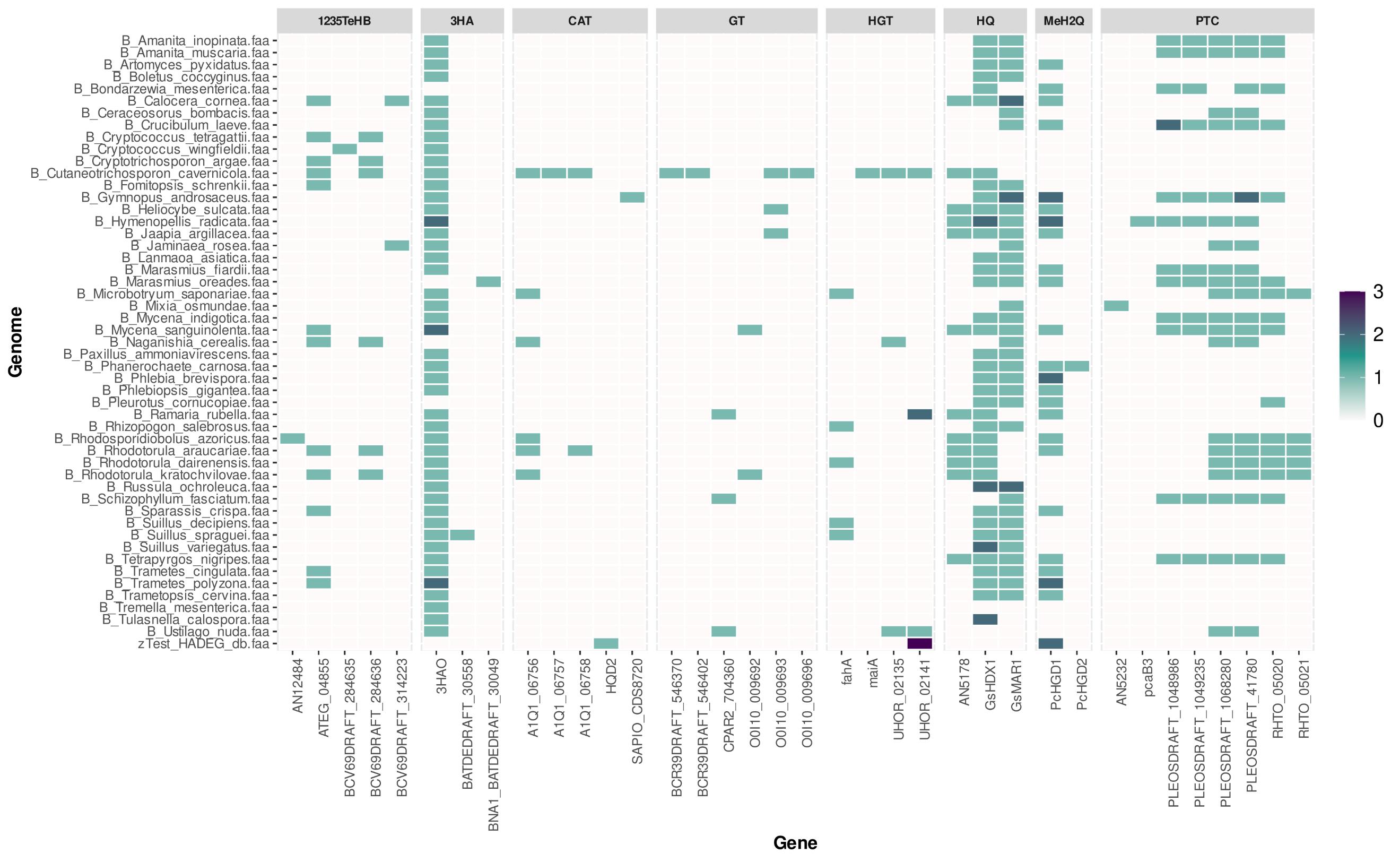

**Figure S5**. FAMeDB analysis of central pathways in fifty random proteomes of Basidiomycota. The number of database hits of the central pathways annotated per proteome and gene (representing an ortholog group) is displayed using a heatmap plot. The last row provides a direct comparison of matches obtained against HADEG database. Results shown consider a minimum sequence identity of 40% in the orthology analysis step. Abbreviations: 1235TeHB - 1,2,3,5‑tetrahydroxybenzene, 3HAO - 3‑hydroxyanthranilate, CAT - catechol, GT - gentisate, HGT - homogentisate, HQ - hydroxyquinol, MeH2Q - methoxyhydroquinone, and PTC – protocatechuate.

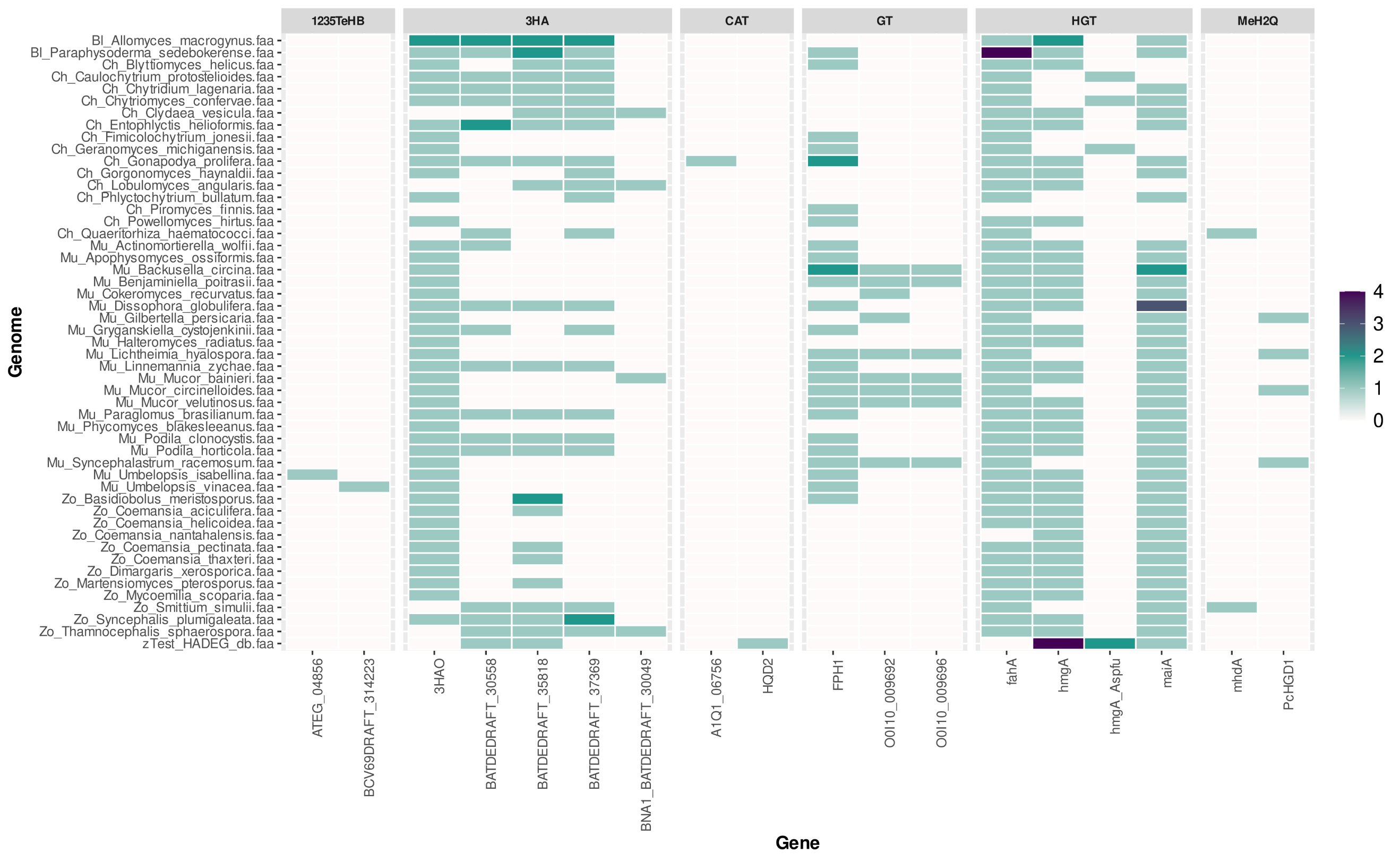

**Figure S6**. FAMeDB analysis of central pathways in fifty random proteomes of non-Dikarya. The number of database hits of the central pathways annotated per proteome and gene (representing an ortholog group) is displayed using a heatmap plot. The last row provides a direct comparison of matches obtained against HADEG database. Results shown consider a minimum sequence identity of 40% in the orthology analysis step. Abbreviations: 1235TeHB - 1,2,3,5‑tetrahydroxybenzene, 3HAO - 3‑hydroxyanthranilate, CAT - catechol, GT - gentisate, HGT - homogentisate, HQ - hydroxyquinol, MeH2Q - methoxyhydroquinone, and PTC – protocatechuate.
